# Cryo-EM structure of native human thyroglobulin

**DOI:** 10.1101/2021.06.06.447243

**Authors:** Ricardo Adaixo, Eva M. Steiner, Ricardo D. Righetto, Alexander Schmidt, Henning Stahlberg, Nicholas M. I. Taylor

**Affiliations:** Center for Cellular Imaging and NanoAnalytics, Biozentrum, University of Basel, Mattenstrasse 26, CH-4058 Basel, Switzerland; Novo Nordisk Foundation Center for Protein Research, University of Copenhagen, Blegdamsvej 3B, DK-2200 Copenhagen, Denmark; Proteomics Core Facility, Biozentrum, University of Basel, Klingelbergstrasse 72, CH-4058 Basel, Switzerland

**Keywords:** thyroglobulin, cryo-electron microscopy, thyroid, hormonogenesis, cryo-electron microscopy, non-homologous insertions, flexible domains

## Abstract

The thyroglobulin (Tg) protein is essential to thyroid hormone synthesis, playing a vital role in the regulation of metabolism, development and growth. Its structure is conserved among vertebrates. Tg is delivered through the secretory pathway of the thyroid follicular unit to the central colloid depository, where it is iodinated at specific tyrosine sites to form mono- or diiodotyrosine, which combine to produce triiodothyronine (T3) and thyroxine (T4), respectively. Synthesis of these hormones depends on the precise 3D structure of Tg, which has remained unknown despite decades of research. Here, we present the cryo-electron microscopy structure of human thyroglobulin (hTg) to a global resolution of 3.2 Å. The structure provides detailed information on the location of the hTg hormonogenic sites and reveals the position as well as the role of many of its glycosylation sites. Our results offer structural insight into thyroid hormonogenesis and provide a fundamental understanding of clinically relevant hTg mutations, which can improve treatment of thyroid diseases.

## Introduction

Thyroglobulin (Tg) is a 660 kDa hyper glycosylated protein expressed in thyrocytes and secreted to the follicular lumen where it accumulates^1^. Dimeric Tg is secreted to the follicular cavity and iodinated to different extents at specific tyrosine residues, a process modulated by the dietary iodine intake^2^. Iodinated Tg is transported to the thyrocyte cytosol by pinocytosis and digested, releasing triiodothyronine (T3) and thyroxine (T4) hormones^3^. Tg is simultaneously a precursor for thyroid hormone (TH) biogenesis and the carrier protein responsible for iodine storage in the follicle colloid. THs are essential to fetal and infant brain development as well as throughout adulthood as metabolism regulators^4^. Mutations in the Tg sequence, or alteration of glycosylation structures, are related to increased risk of thyroid cancer as well as dyshormogenesis associated with goiter^5–7^.

Analysis of primary sequences of Tg allowed early identification of internal homology domains classified as type 1, type 2 and type 3 repeats^8^, as well as a cholinesterase-like domain (ChEL) at the carboxyl end of the protein^9^. Type 1 repeats occur 11 times in the human Tg (hTg) sequence and are homologous to 1a domains found in other proteins with known structure^10,11^. The ChEL domain also has several homologues with structures determined by X-ray diffraction^12–15^. The ChEL domain assists Tg folding, dimerization and secretion processes^16^.

Despite the extensive biochemical characterization of Tg in the past decades^1,17–22^ the three-dimensional structure of Tg remained unknown^1^, limiting the understanding of its function. Here, we present the structure of endogenous hTg, determined by cryogenic transmission electron microscopy (cryo-EM). The obtained EM map depicts a dimer with extensive interchain contacts which include but are significantly larger than the ChEL dimer interface. The hTg monomer has 57 disulfide bridges (DSB), which add structural stability and rigidity to most of the protein. However, the extreme N- and C-terminal segments as well as two other regions (the so-called “foot” and “wing”) display a higher degree of flexibility, which is likely related to function.

We provide a comprehensive structural description of the endogenous hTg dimer and demonstrate the functional importance of the natural post-translational modification and iodination sites by presenting an atomic model of the nearly complete protein. The native hTg sample is heterogeneous both in composition and conformational states, which likely represents the *in vivo* requirements for hTG function.

While this work was in preparation, Coscia et al.^23^ described the structure of native, deglycosylated hTG as well as recombinant, non-deglycosylated hTG, identified the putative hormonogenic sites and validated them using an *in vitro* hormone production assay. Our findings are consistent with those of Coscia et al. but our cryo-EM reconstruction is of the native, non-deglycosylated hTG and of relatively better resolution. We additionally describe novel posttranslational modifications of hTG.

## Results

We obtained a homogenous solution of hTg from native sources, by resuspension and gel filtration of the commercially available lyophilate without performing any in vitro iodination. The cryo-EM images of plunge-vitrified hTg solution displayed monodisperse, randomly oriented particles with size and shape consistent with previous observations in negative stain preparations^17^. A cryo-EM map at 3.2 Å nominal resolution was reconstructed allowing atomic modelling of the hTg dimer to around 90% completeness (2,483 modelled residues over 2,748 expected residues per chain). This composite map is the result of a globally-refined consensus map and two maps locally refined around particularly flexible regions, the “wing” and the “foot”, as described below (**Fig. 1, Supplementary Video 1**). These peripheral and flexible domains are likely an obstacle to obtain diffraction-quality crystals, which probably prevented the structural determination of hTg in the past. The hTg dimer is approximately 250 Å long by 160 Å wide and 110 Å along the C2 symmetry axis. Each chain is formed by regions I, II and III and a C-terminal ChEL domain (**Fig. 1**). The interface between monomers buries an area of 31,100 Å^2^ involving all regions except region II. The hTg structure is annotated similarly to what is reported in the literature^1^, however, region I lacks the so called “linker” between repeats 1.4 and 1.5. Three types of cysteine-rich internal homology repeats are present in hTg: type 1, type 2 and type 3. There are 10 type 1 repeats within region I and an 11th in region II. Three type 2 repeats, each bearing 14 to 17 residues, lie between the hinge region and repeat 1.11. Type 3 repeats are located between repeat 1.11 and the ChEL domain.

**Figure 1.**
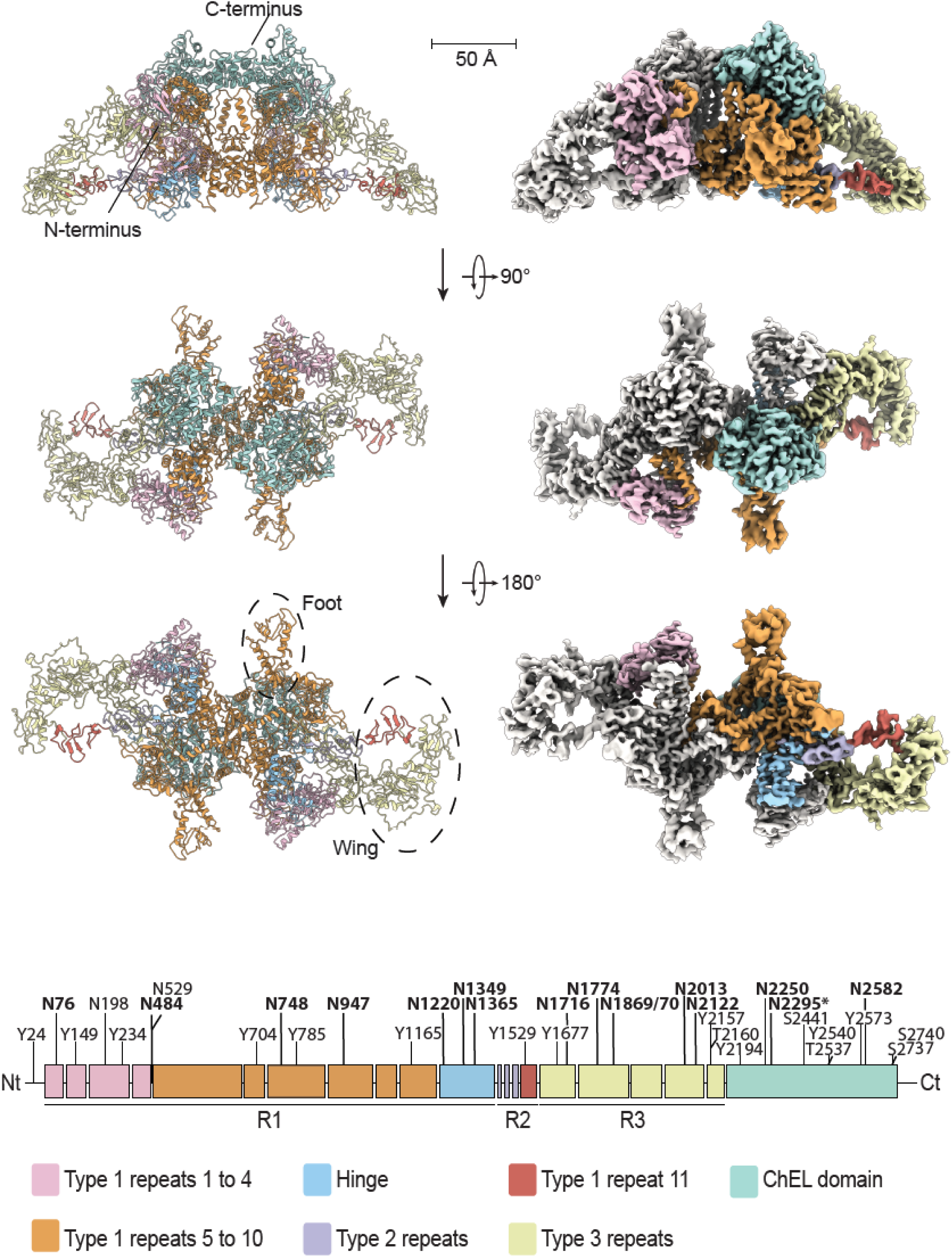
The cryo-EM map of hTg at 3.2 Å. (right) Density map of hTg with one monomer in white and corresponding atomic model (left). Map and model are colored as in the bottom linear diagram of a single hTg monomer.

### Type 1 repeats

The first four type 1 repeats in the proximal region I cluster at the N-terminus of each chain and establish extensive contacts with region III and the hinge of the opposing chain, as well as intra chain contacts with repeat 1.5 (**Fig. 1, 2**). Within repeats 1.1 to 1.4 we observe two non-homologous insertions (NHI), namely on loops 2 and 3 of repeat 1.3. Repeats 1.5 to 1.10 occupy the central core of hTg and this ensemble forms contacts to all other regions on both chains. Additional NHIs are present in loop 1 of repeat 1.5, loop 2 of repeat 1.7 and loop 2 of repeat 1.8 (**Fig. S6**). Insertions of repeats 1.3 and 1.5 are in close proximity and exposed between the proximal region I and the ChEL domain of the opposing chain. Insertions of repeat 1.8 from both chains lie at the C2 symmetry axis and form a helix bundle providing additional 1,710 Å^2^ interchain contact surface (**Fig. 3**). Repeat 1.7 exposes an NHI protruding almost radially to the C2 symmetry axis and forms no additional contacts. We named this protrusion as “foot”, and it is flexible as suggested by the diffuse density obtained in the consensus map. Residues 378 to 615 were assigned to the so called “linker region” in previous work while in our structure they form the insertion of repeat 1.5. A consequence of our annotation is that repeat 1.5 encompasses 2 additional cysteines, Cys408 and Cys608, and a total of 4 disulfide bridges, therefore we classify this repeat as type 1c, as opposed to type 1a and type 1b repeats containing 6 and 4 cysteine residues respectively. The remaining NHIs are devoid of additional cysteine residues.

**Figure 2.**
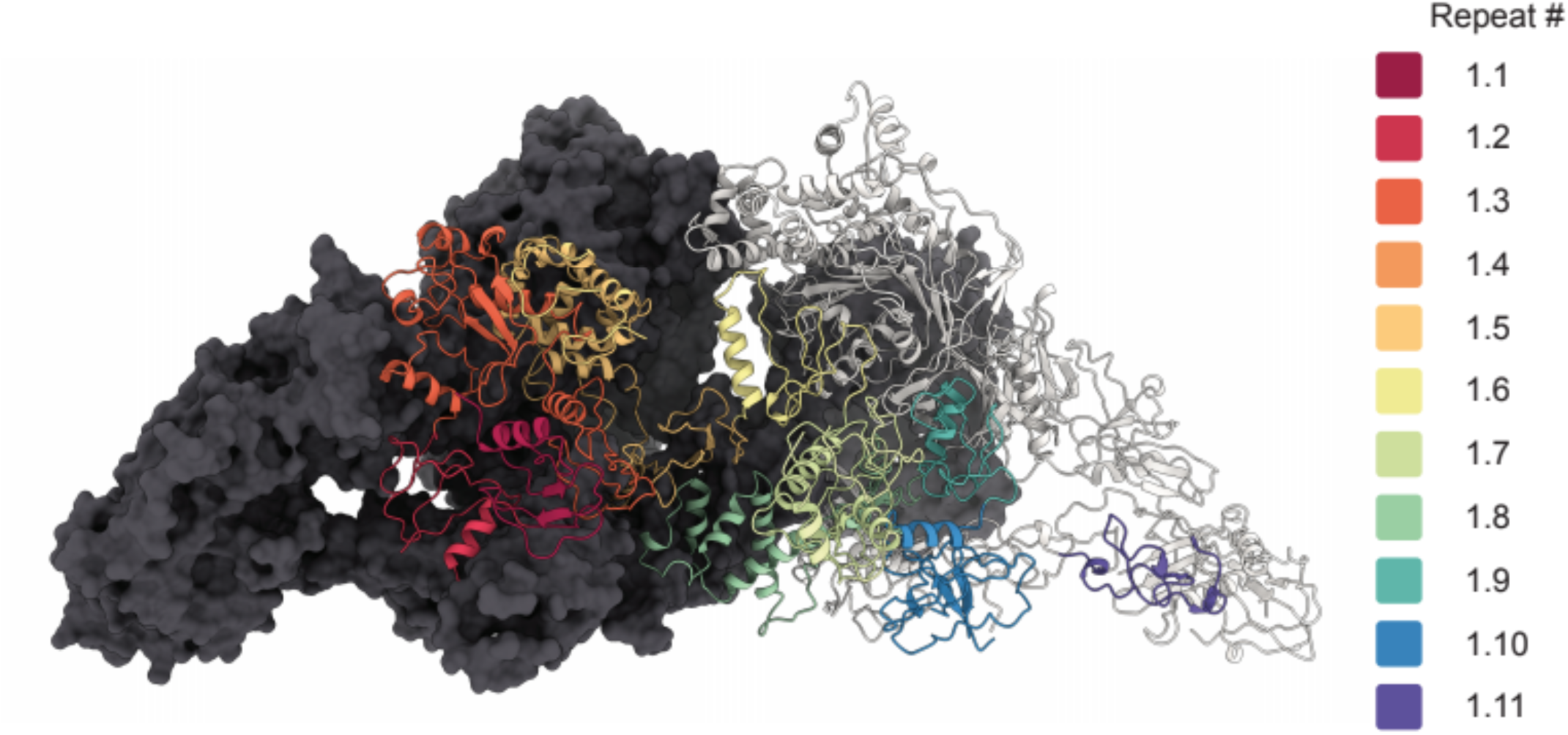
Disposition of the different type 1 repeats in the hTg structure. Surface representation of one hTg monomer in the background colored charcoal. Ribbon representation of the second hTg monomer in the foreground with each type 1 repeat colored as in the scheme on the right.

**Figure 3.**
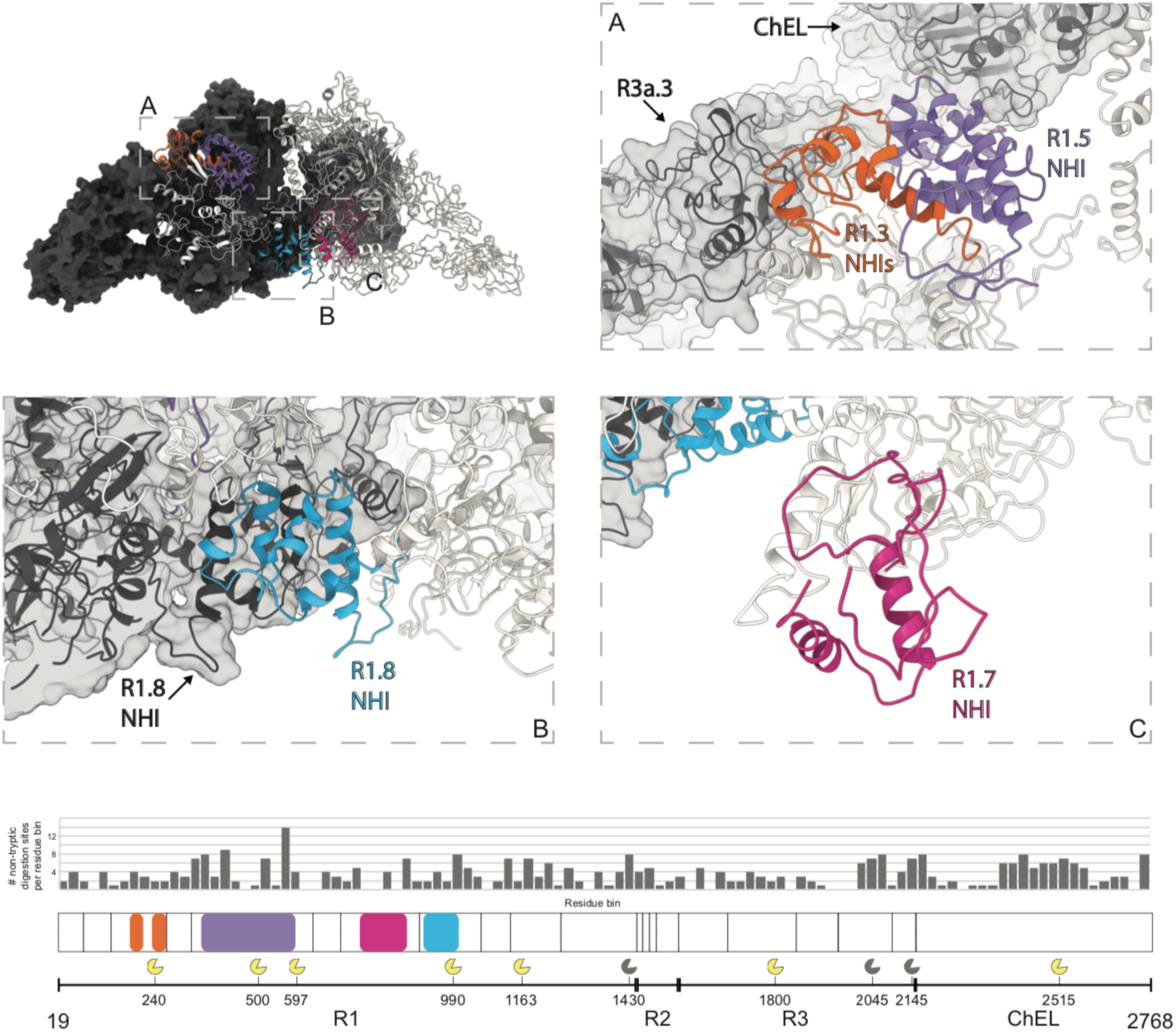
Location of type 1 repeat NHIs and proteolysis clusters. (Top) Surface representation of one hTg monomer (in the background) colored charcoal. Ribbon representation of the second hTg monomer in the foreground with each NHI colored differently: orange – repeat 1.3 NHIs; purple – repeat 1.5 NHI; blue - repeat 1.8 NHI; wine – repeat 1.7 NHI. Detailed clipped views of each insertion are displayed in the dashed boxes; box B view direction is the same as the top left image while boxes A and C were reoriented for better depiction. (Bottom) Histogram of the non-tryptic cleavage sites detected by MS with major sectors depicting the approximate position of previously reported (yellow) and novel (grey) cleavage clusters.

### Type 2 repeats

hTg residues 1456 to 1487 comprise three contiguous type 2 repeats flanked by the hinge and repeat 1.11. Each type 2 repeat comprises 2 cysteine residues, all of which are engaged in disulfide bond formation (**Fig. 4**). The most N-terminal cysteine of each type 2 repeat establishes a disulfide bond (DSB) with the adjacent N terminal domain while the most C-terminal cysteine establishes a DSB with the adjacent C-terminal domain. Therefore, both repeat 1.11 and hinge region are linked to repeats 2.1 and 2.3 via DSBs contrary to previous hypothesis^8^ where all 6 cysteines in type 2 repeats would form DSBs internally.

**Figure 4.**
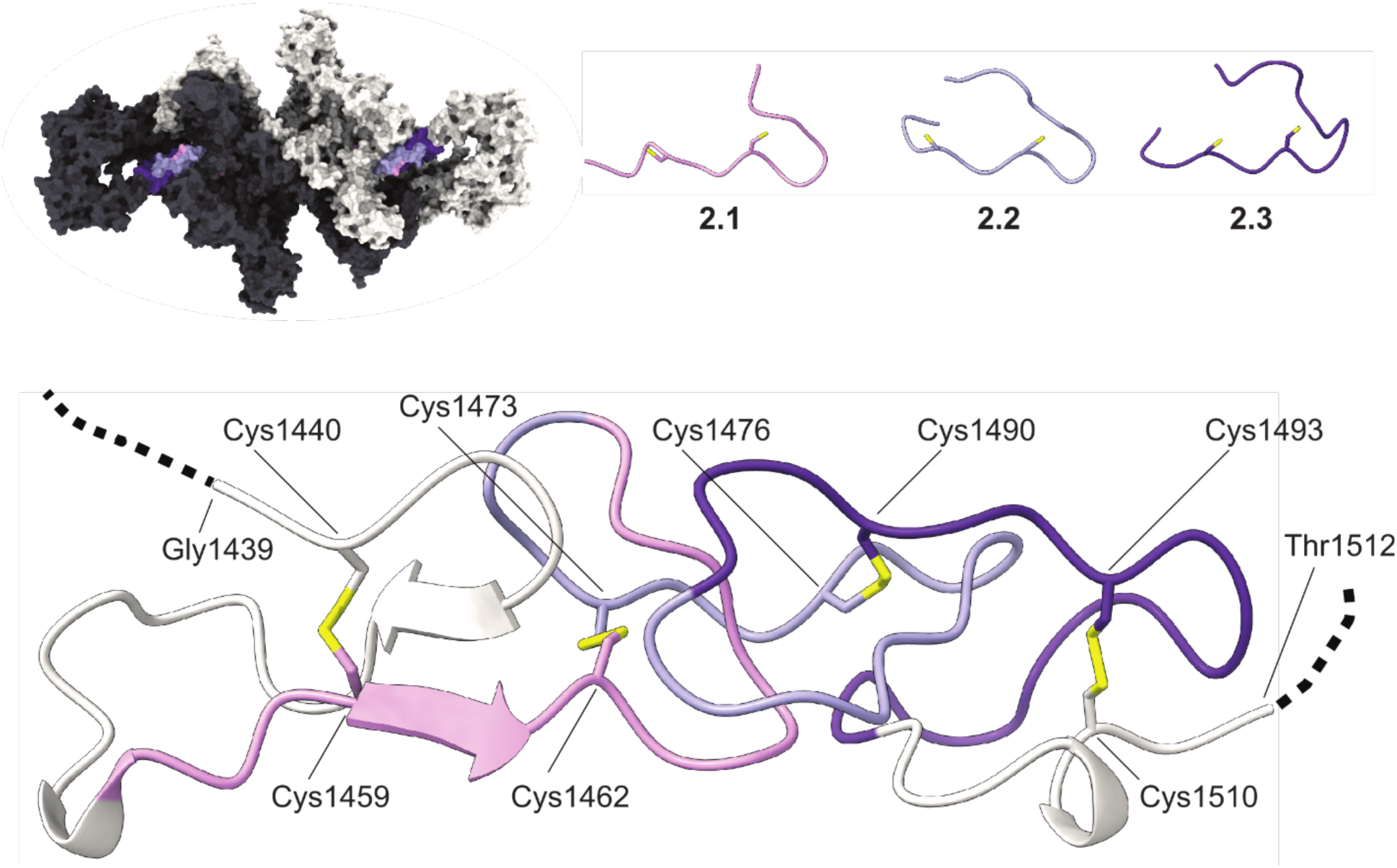
Type 2 repeats. (Top) Ribbon representation of the aligned type 2 repeats. (Bottom) Ribbon representation of the type 2 repeats as disposed in the context of the hTg structure. Cysteine residues represented in sticks.

Repeat 2.1 has a small beta-strand between both cysteine residues which is the only secondary structure element found within the set. All type 2 repeats have a shape reminiscent of an arrowhead where the pointy edge of one repeat is embraced by the flat base of the following type 2 repeat.

It has been hypothesized that the CXXC motif in type 2 repeats constitutes a thioredoxin box which may be required for Tg multimerization via intermolecular DSBs^24^. We did not investigate this hypothesis. However, all type 2 repeats are extensively solvent exposed and therefore could serve as a potential substrate to thioredoxin.

### Type 3 repeats

Region III comprises a total of 5 type 3 repeats linking the ChEL domain to repeat 1.11. These repeats are formed by an alpha helix followed by a three-stranded beta sheet and can be subdivided into type 3a, bearing 8 cysteines, and type 3b, bearing 6 cysteines (**Fig. 5**). The loop connecting the third beta strand to the neighbor domain is longer and apparently disordered. We noticed the previous annotation^1^ places the limits of each type 3 repeat within secondary structure elements and therefore does not take into consideration the globular nature of individual domains which can be discerned in the structure but, for the sake of consistency, we follow the same annotation.

**Figure 5.**
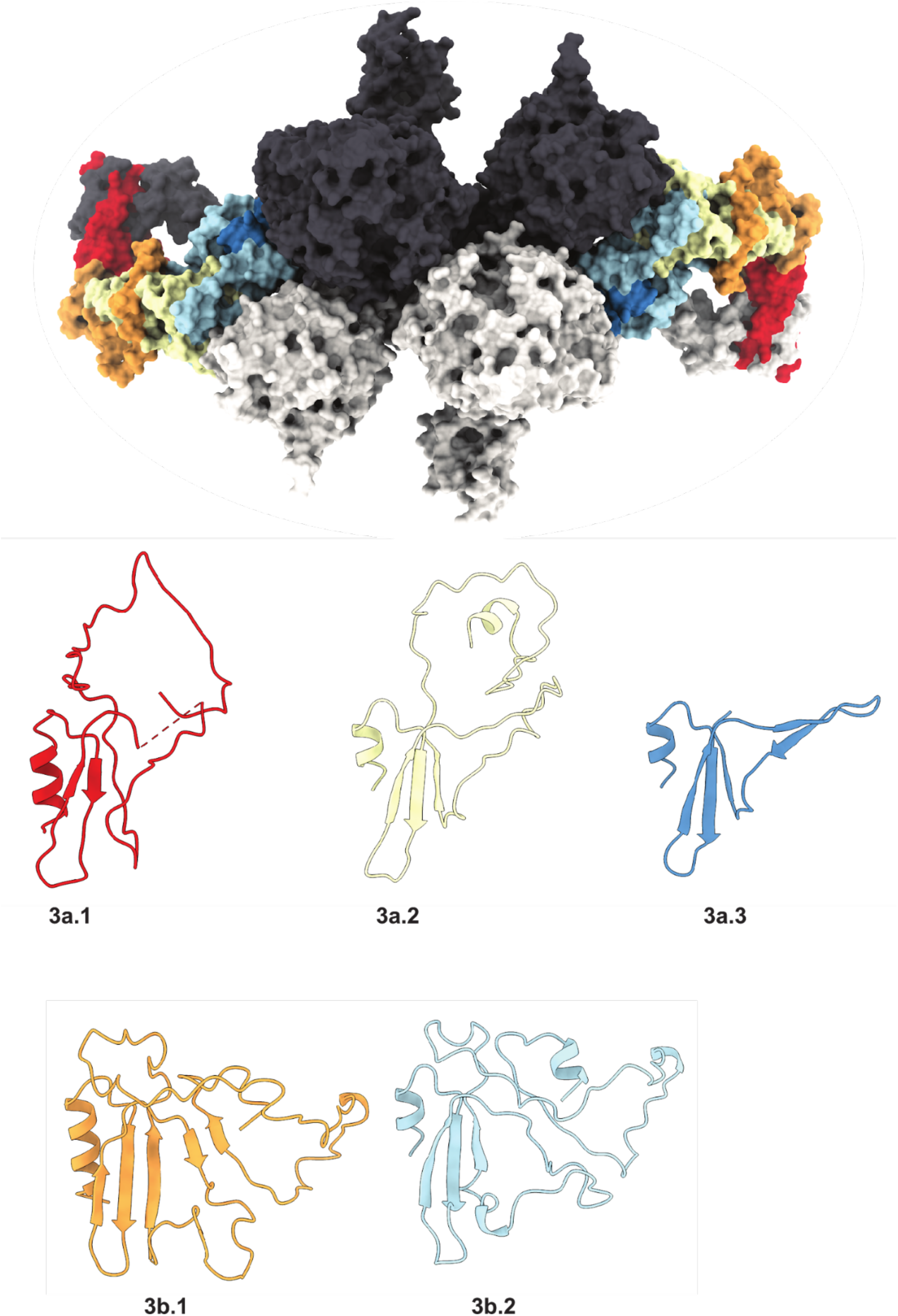
Type 3 repeats. Ribbon representation of the aligned type 3 repeats.

Repeats 3a3 and 3b2 were reasonably well defined in the consensus map (see **Materials and Methods**) however this was not the case for repeats 3a2, 3b1, 3a1 and down to repeat 1.11. Both 2D classes and consensus refinement maps were not well defined in the 3a2 to 1.11 region (which we named “wing region”) likely due to increased flexibility. The wing has a “C” shape where the tips seem to act as pivot points, connecting repeats 2.3 to 1.11 at the N-terminus and repeats 3a2 to 3b2 at the C-terminus. The reason for the increased flexibility of the wing region is unknown to us.

### Mapping of hTg’s hormonogenic sites

The post translational iodination of hTG contributes both to thyroid hormonogenesis as well as iodine storage. One hTg monomer contains 66 tyrosines, therefore the hTg dimer represents a huge reservoir for iodination and post-translational modifications. Under sufficient iodide intake 10-15 tyrosine residues become mono- and diiodotyrosines (MIT, DIT), serving as functional hormonogenic units within the hTg structure^3^. Hormonogenesis requires a selected pair of donor and acceptor tyrosine residues and four of such sites (A-D sites) were proposed in hTg^21,25^ (**Table 1**). Coupling is proposed between a DIT and another DIT (donor and acceptor) or between a MIT and a DIT (MIT donor and DIT acceptor) to undergo an oxidative quinol-ether coupling reaction to form T4 or T3, respectively^20^.

**Table 1:**
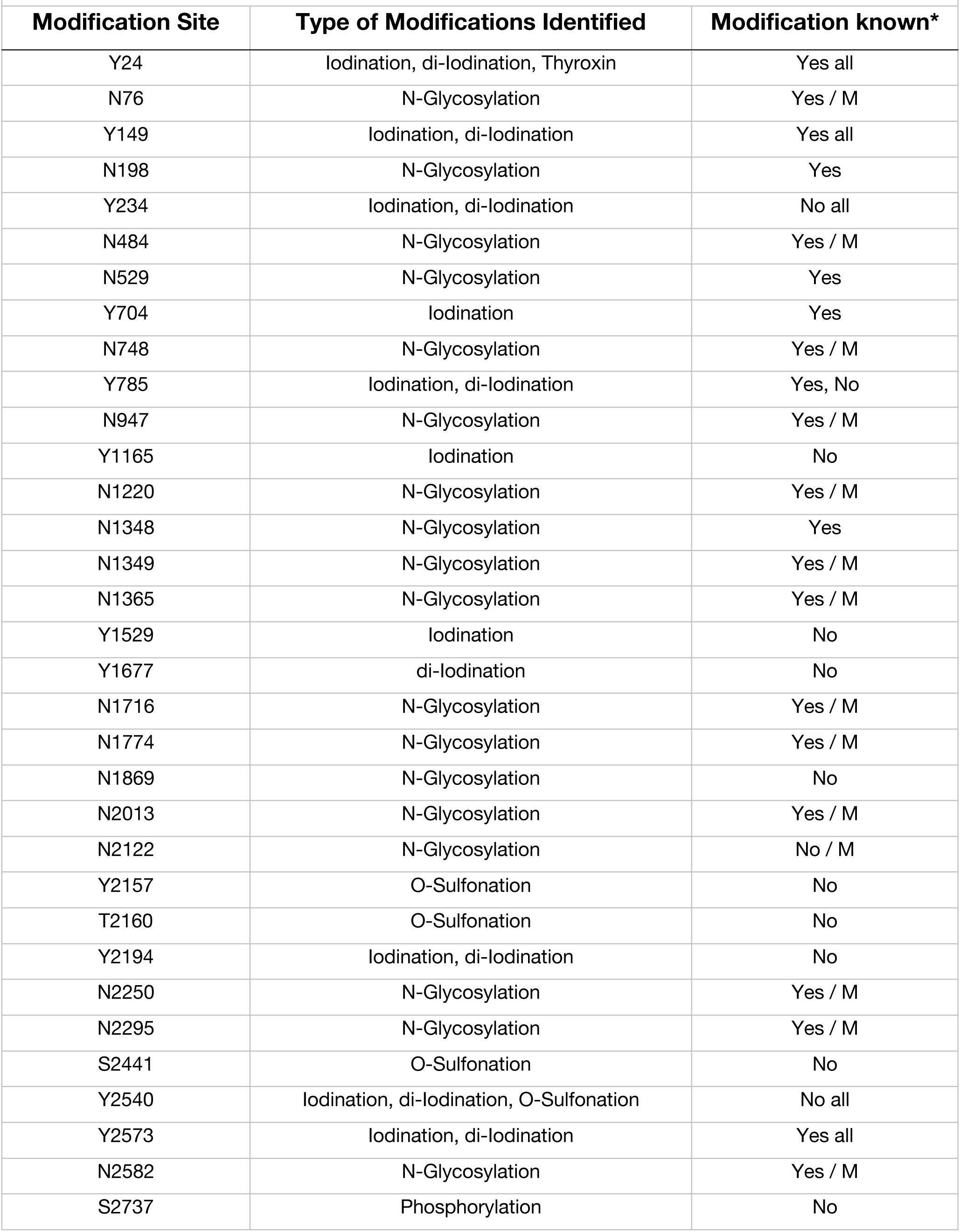
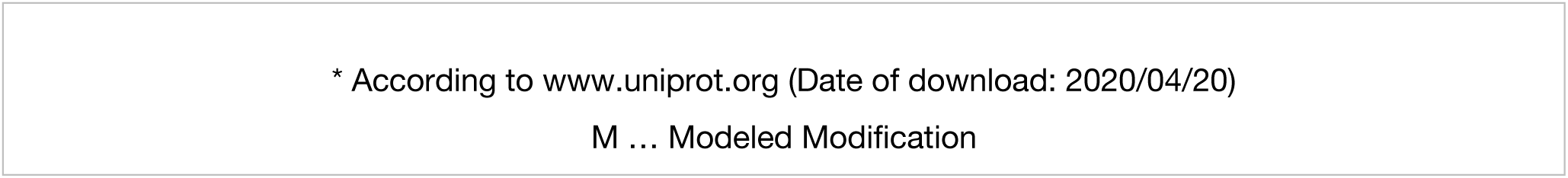
List of Native Human Thyroglobulin Modifications Identified by LC-MS.

| **Modification Site** | **Type of Modifications Identified** | **Modification known\*** |
| --- | --- | --- |
| Y24 | Iodination, di-iodination, Thyroxin | Yes all |
| N76 | N-Glycosylation | Yes / M |
| Y149 | Iodination, di-iodination | Yes all |
| N198 | N-Glycosylation | Yes |
| Y234 | Iodination, di-iodination | No all |
| N484 | N-Glycosylation | Yes / M |
| N529 | N-Glycosylation | Yes |
| Y704 | Iodination | Yes |
| N748 | N-Glycosylation | Yes / M |
| Y785 | Iodination, di-iodination | Yes, No |
| N947 | N-Glycosylation | Yes / M |
| Y1165 | Iodination | No |
| N1220 | N-Glycosylation | Yes / M |
| N1348 | N-Glycosylation | Yes |
| N1349 | N-Glycosylation | Yes / M |
| N1365 | N-Glycosylation | Yes / M |
| Y1529 | Iodination | No |
| Y1677 | di-iodination | No |
| N1716 | N-Glycosylation | Yes / M |
| N1774 | N-Glycosylation | Yes / M |
| N1869 | N-Glycosylation | No |
| N2013 | N-Glycosylation | Yes / M |
| N2122 | N-Glycosylation | No / M |
| Y2157 | O-Sulfonation | No |
| T2160 | O-Sulfonation | No |
| Y2194 | Iodination, di-iodination | No |
| N2250 | N-Glycosylation | Yes / M |
| N2295 | N-Glycosylation | Yes / M |
| S2441 | O-Sulfonation | No |
| Y2540 | Iodination, di-iodination, O-Sulfonation | No all |
| Y2573 | Iodination, di-iodination | Yes all |
| N2582 | N-Glycosylation | Yes / M |
| S2737 | Phosphorylation | No |

\* According to www.uniprot.org (Date of download: 2020/04/20)
M ... Modeled Modification

To map the iodination status and positions of hTg, we used mass spectrometry (MS) and detected 11 iodinated tyrosine residues that could serve as potential donor and/or acceptor site for TSH production (**Fig. 1, Table 1**). Out of the detected sites, 10 were MIT, 6 sites MIT and DIT, 3 only MIT and 1 site DIT only. A special role, as highlighted before, plays Y24, the most efficient T4 forming unit, where we detected a thyroxin T4 modification as well^21^ (**Table 1**).

We did not observe convincing densities for iodine in the cryo-EM map, probably because iodination levels were low, these sites may be particularly sensitive to radiation damage during data collection, the iodination pattern may be inconsistent between different particles, or a combinations of all these factors.

Despite the large number of iodination sites, hormonogenesis depends primarily on tyrosine residues near the N-terminus (the A-site with acceptor at Y24, crucial for T4 synthesis) and the C-terminus (the C-site with acceptor at Y2766, specific for T4 and T3 synthesis) respectively. Further proposed hormonogenic sites are site B, with acceptor Y2573^18^(iodination detected by MS), site D (acceptor Y131)^25^ (no iodination detected by MS) and site E (acceptor Y704)^18^(MIT iodination detected and modeled) (**Fig. 6, Table 1**).

**Figure 6.**
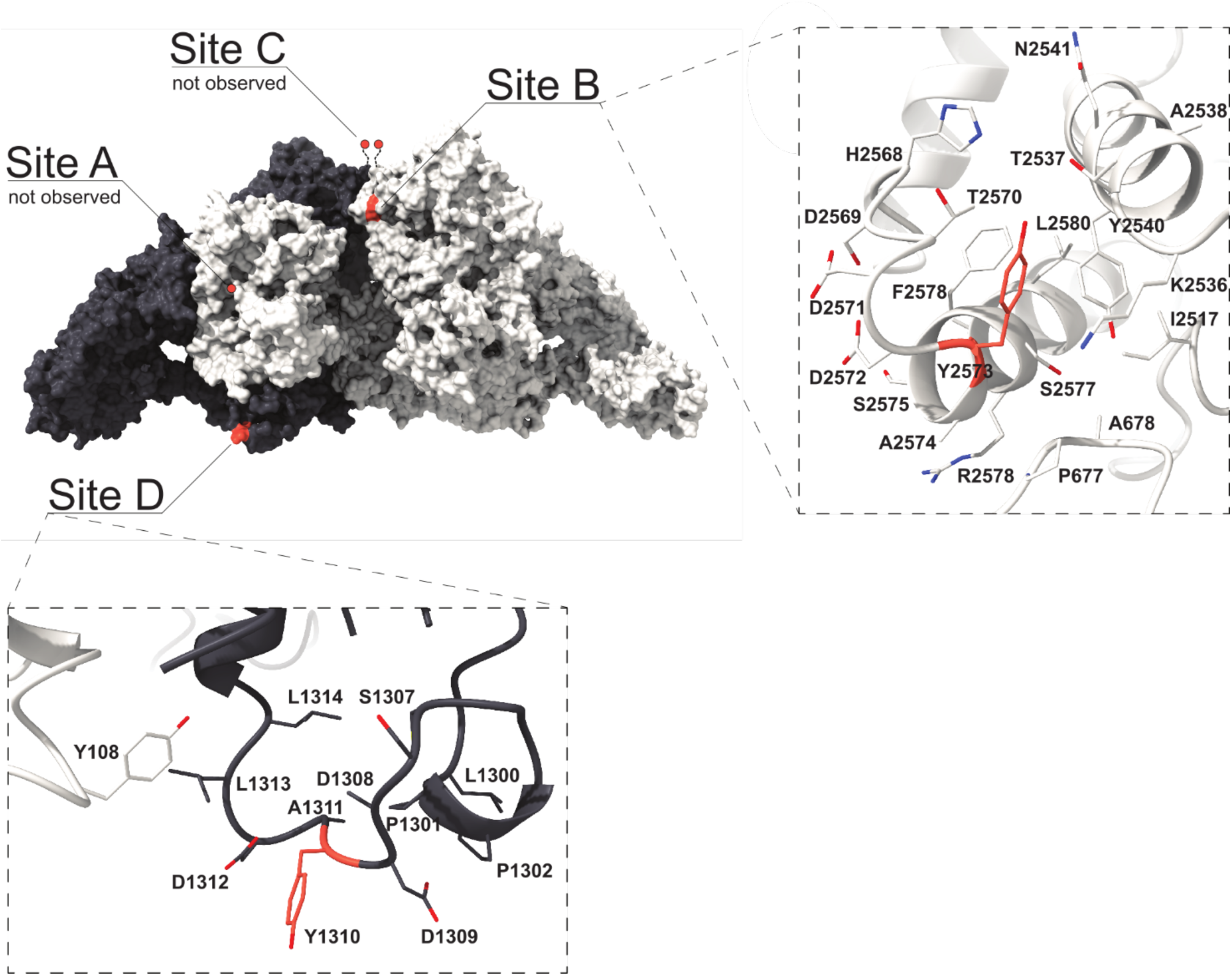
Location of hormonogenic sites in hTg. (Top Left) hTg surface representation with one monomer in charcoal and one monomer in white. Acceptor tyrosine residues in each site (A to D) are marked in orange.

In case of the A-site, although we do not directly observe the acceptor Y24 in the map we can deduce its approximate location as the first modeled residue is P30. Furthermore, we revealed Y24 to be clearly mono- and diiodinated and as well showing an additional T4 mass (**Table 1**). Potential donor sites for Y24 have been suggested to be Y234 (donor 1) or Y149 (donor 2)^25^. We detected mono and di-iodination for both donor sites Y149 and Y234 providing further evidence supporting that these residues are potential donors within hormonogenic site A^25^. Taken together with the distance to P30 and flexibility of the N-terminal, this suggests that T4 synthesis can occur within a single hTg monomer.

Tyr258 (which is also close to a glycosylation site N484) has been suggested as an alternative donor. In the structure, this residue is still in relatively close position to Pro30 and therefore Y24, however, major conformational changes would be necessary given that the sidechain of Y258 is oriented towards the hTg core. At the B-site, Y2573 is located at the surface of the protein and fairly accessible. The residue is proposed to be an acceptor whereas Y2540 functions as donor (**Fig. 6**). For both donor and acceptor tyrosines, we observed mono and di-iodination by mass spectrometry and the 6 Å close contact would allow a coupling reaction. In 15 Å proximity is as well Y2478 but has not been shown to be iodinated in our sample. Interestingly, T2537, at 3.8Å and 5.6 Å distance to both tyrosines, Y2573 and Y2540, at the B-site, was found to be phosphorylated therefore maybe playing a crucial role in acceleration of hormone production^26^. At about 28 Å distance from this T3 production site, S2441 was detected by mass spectrometry to be sulfonated (**Table 1**). Sulfonated serines can be involved in various functions including protein assembly and signaling processes and this type of PTM was detected in a cathepsin-C like enzyme from parasites^27^.

The C-site is not visible in the structure, where T2727 is the last residue traced. It has been shown that Y2766 is acceptor to the Y2766 in the neighboring monomer of the hTG dimer^28^. Our structure is perfectly consistent with this observation: the C-α distance between the Thr2708 residues of both monomers is only 20 Å.

The D-site, with a captor Y1310, is accessible and exposed. The donor residue has been proposed to be Y108^25^. We did not observe iodination of Y1310 nor Y108 and our data does not support those residues as a D-site *in vivo*. However, since those residues are surface exposed, *in vitro* iodination was shown to be possible^25^.

A putative additional site, located around Y704, could also be found in the structure, as it is accessible at the surface of the protein and had monoiodination detected (**Table 1**). Several donor or acceptor sites are present: Y866 (26 Å distance to Y704), Y883 (20 Å), Y2640 (14 Å), Y2637 (19 Å), but none were detected to be iodinated. However, Y866 and Y883 were found to be iodinated *in vitro* and could therefore be potential donors^18^.

A wide variety of other iodinated sites (Y785 MIT/DIT, Y1165 MIT, Y1529 MIT, Y2194 MIT/DIT) have been identified and most of these are located on the surface (**Table 1**). However, it is unclear to what extent they have a role in hormonogenesis as opposed to iodine storage.

### Mapping of proteolysis sites

The lifecycle of Tg comprises multiple proteolysis events: the cleavage of the N-terminal 19 residues signal peptide; the N- and C-terminal cleavages which liberate iodopeptides; the limited proteolysis of Tg which releases Tg particles from the colloid agglomerate; and, finally, the digestion of Tg internalized in the thyrocyte^29–31^. Cathepsins, a family of cysteine proteases, perform the mentioned proteolytic attacks on Tg both inside the thyrocyte and in the follicular lumen. The approximate locations of cathepsin proteolysis sites are depicted in **Fig. 3**. Digestion of the extreme N- and C-terminus containing thyroid hormones (TH) is among the earliest proteolysis events experienced by mature Tg^32–34^.

Insertions of repeats 1.3, 1.5 and 1.8 contain motifs targeted by cathepsins and a sequence of proteolysis events has been suggested^35^ involving different proteases from the cathepsin family. Hence, one function of these insertions is to expose proteolysis-prone segments in order to facilitate hTg digestion.

Our hTg atomic model lacks the residues between N496-P547 and T1781-N1814 because no clear density was observed in those regions in the EM map. Interestingly, two of the proteolysis sites plotted in **Fig. 3** (major sector marks on positions 500 and 1800) lie within these missing segments, suggesting that our sample was partially digested at these specific locations.

### hTG glycosylation and other modifications

The addition of glycan structures to hTg is crucial for protein folding, structure and therefore function, immune-recognition, cell signaling and play a significant role in protein transport and THS production^36^. Within the human thyroglobulin monomer 16 N-linked glycosylation sites have been discovered in the mature protein^19,25^.

We identified 16 N-linked glycosylation sites, per hTG monomer, by mass spectrometry (**Fig. 1, Table 1**). Out of those, 14 asparagine residues showed an additional density in the map and N-linked glycan structures were modeled. The two glycosylation sites, N198 and N529, are located close to the hormonogenic A-site and within the region of a proteolytic site, respectively and therefore not modelled (**Fig. 1**). Based on the map an additional glycosylation site at position N2295 was identified and a high mannose was modeled indicating that there might be more than 16 sites in the mature hTg protein (**Table 1**)^25^.

We present 3 new glycosylation sites at position N198, N1869 and N2122, that have not been described in previous biochemical studies (**Table 1**)^19^. For several of the previously annotated glycosylation sites we did not detect any modification nor did we observe any additional map density; N110, N198, N816 and N1348^19,25^. Besides glycosylation, we have also mapped sulfonation, phosphorylation and acetylation. We identified 15 acetylation sites, 4 phosphorylation and 4 sulfonation sites (**Table 1**). No methylations or succinations were detected.

The phosphorylation at T2537 (PO_4_) and sulfonation Y2540 (SO_3_) are within the hormonogenic B-site. Phosphorylation is thought to improve the efficiency of T3 formation^26^ and sulfonation of Y24 and the surrounding peptide sequence was shown to be crucial in thyroid hormone synthesis^37,38^.

Sulfated iodotyrosines (Tyr-S) have a short life before the coupling reaction occurs and it is suggested that after Tyr-S binding to peroxidase where it is iodinated, the sulfate group is removed, releasing an iodophenoxy anion available for coupling with an iodotyrosine donor^38^. For Y2540, the donor residue in the B-site, we detected sulfonation, MIT an DIT representing all 3 states of hormone site preparation and a perfectly prepared hormone formation site (**Fig. 6**).

### Mapping of nonsense and missense mutations

We considered all the nonsense and missense mutations reported previously^1^ and plotted these on the hTg structure (**Fig. 7**). The majority of the mutations causing early termination of hTg translation as well as those causing a change in amino acid identity fall within modelled regions; 31 of 36 nonsense mutations and 84 of 90 missense mutations. Interestingly, 4 of the 5 major clusters of the mentioned mutations (**Fig. 7**) overlap with the proteolysis sites depicted in **Fig. 3**, namely those within repeats 1.3, 1.8, 1.10 and to lesser extent the ChEL domain.

**Figure 7.**
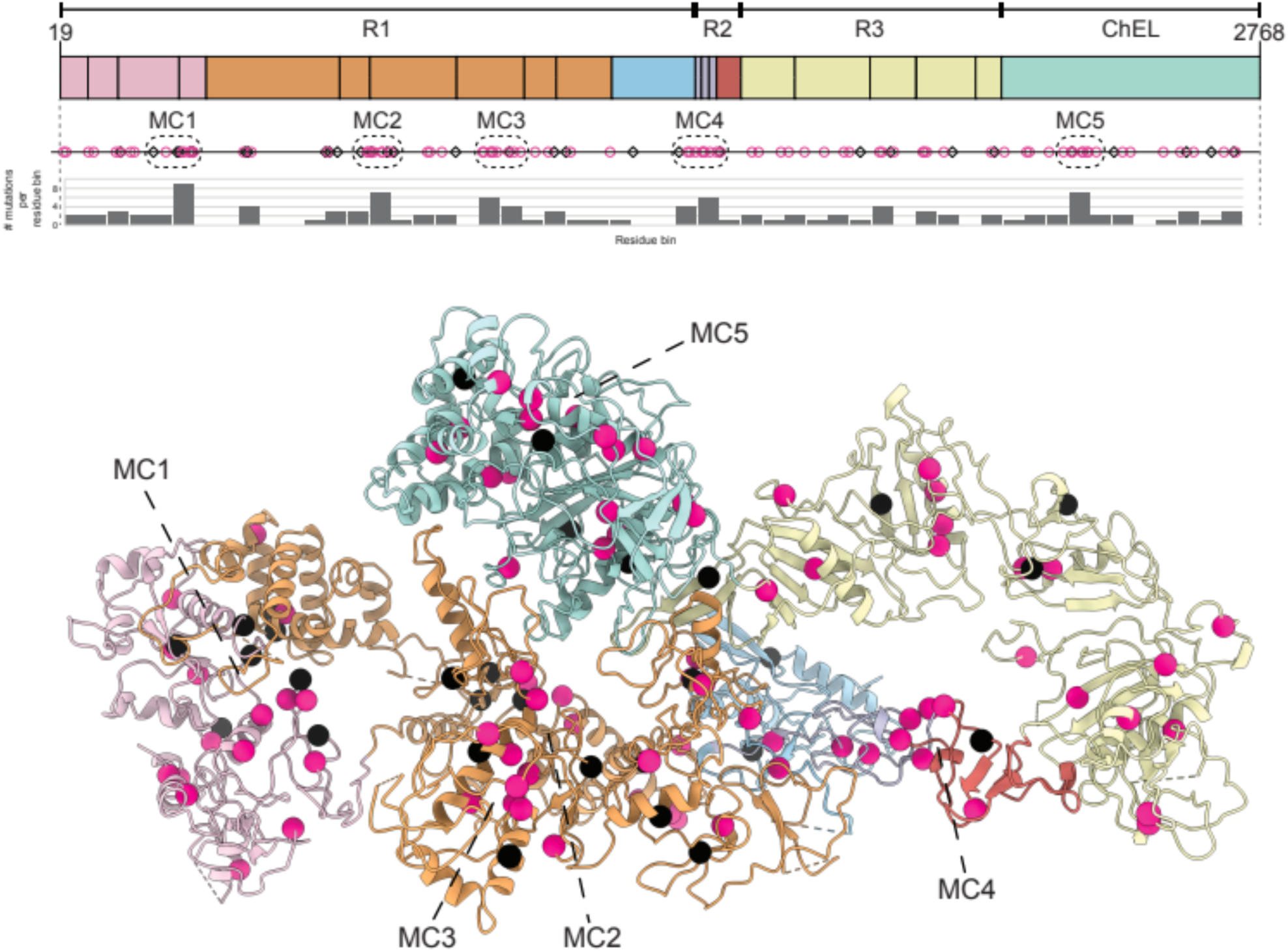
Location of nonsense and missense mutations in hTg. (Top) hTg linear diagram depicting the location of nonsense mutations in black squares and missense mutations in pink; domains color code is the same as **Fig. 1.** Dashed boxes represent the 5 mutation clusters (MC1 to MC5) with the highest density of mutations. (Bottom) Location of the same mutations in the ribbon representation of hTg.

## Discussion

We determined the atomic structure of hTg based on a composite cryo-EM density map at an overall resolution of 3.2 Å for the consensus map. The overall dimeric structure is consistent with previous biochemical experiments, which showed that the cholinesterase domain is necessary and sufficient for hTg dimerization, at least when overexpressed in HEK293 cells^39^, but also reveals the participation of regions 1 and 3 in dimer contact formation.

The ChEL domains and repeats 1.6 to 1.8 form the core of the dimer and lie in close proximity to the C2 symmetry axis while regions 2 and 3 as well as the remaining type 1 repeats occupy more peripheral zones. We modelled 2 of the expected 4 hormonogenic sites, namely site B and site C. Acceptor tyrosines of sites A and C are located at the extreme N and C terminus of the hTg chain and could not be modelled likely due to inherent flexibility of these regions.

Our structural characterization also evidences features of hTg that were previously unknown: the environment and possible function of the 4 NHIs present in type 1 repeats; the previously annotated linker region which in fact is a NHI of repeat 1.5; the globular nature of type 3 repeats, which could be annotated differently, and finally the flexible nature of the wing and foot regions.

All NHIs are solvent exposed and present peptide motifs recognized by proteases as determined by MS analysis. The NHI of repeat 1.7 is unique in the sense that the remaining NHIs establish contacts to adjacent domains other than the type 1 repeat itself. One speculative hypothesis is that repeat 1.7 NHI could still form inter domain contacts but in the context of hTg multimerization.

The hTg atomic structure is decorated with a variety of post translational modifications which were further studied by MS. Importantly, we describe three new glycosylation sites. The density of detected proteolysis sites and the multiple iodination states found for Y24 strongly indicate that our hTg sample was heterogeneous.

We expect the hereby-presented structure of native hTG leads to an improved understanding of Tg biology that could be applied in the diagnosis and therapy of thyroid disease, where our model could be valuable e.g. to determine the location of different antibody epitopes and their relation to autoimmune diseases.

## Methods

### Sample preparation

Human Thyroglobulin (catalog no. T6830; Sigma-Aldrich) was dissolved in gel filtration buffer (25 mM Tris-HCl pH 7.5, 150 mM NaCl, 1x sodium azide) and injected onto a Superdex-200 increase size-exclusion chromatography column connected to an ÄKTA purifier FPLC apparatus (GE Healthcare Bio-Sciences). Peak fractions were pooled and concentrated to 2 mg/mL of protein before plunge freezing.

### Cryo-EM sample preparation and data collection

Quantifoil 2/2 400 mesh Cu grids were glow discharged in low pressure air for 30 s. Then 3 μL of concentrated hTg were dispensed to the hydrophilic surface of the grid prior to single side blotting for 2 s and plunge freezing in liquid ethane using a Leica EM GP2 plunger (Leica Microsystems) operating at 20 °C and 80% relative humidity.

Frozen grids were imaged using a Titan Krios (Thermo Fisher Scientific) transmission electron microscope operating at 300 kV equipped with a Gatan Quantum-LS energy filter (slit width 20 eV; Gatan Inc.) and a K2 Summit direct electron detector (Gatan Inc.). SerialEM^40^ was used for automated data collection with 7 acquisitions per hole using beam-image shift^41^. Movies were recorded in counting mode with a pixel size of 0.64 Å/px at the sample level. Each movie comprised an exposure of 50 e^−^/Å^2^ fractionated into 50 frames over 10 s.

### Image processing and model building

Movies were preprocessed online in FOCUS^42^ using MotionCor2^43^ for drift correction and dose weighting and CTFFIND 4.1^44^ for contrast transfer function estimation. Out of 8,119 movies acquired, 4,504 had an estimated CTF resolution better than 4 Å and were selected and used for automated particle picking in Gautomatch (Zhang, K., https://www.mrc-lmb.cam.ac.uk/kzhang/Gautomatch/) with a CC threshold of 0.4 using a Gaussian blob as template.

Particles were classified in 2D using RELION-3^45^ and the best classes were selected for ab-initio map generation and auto-refinement. EMAN2^46^ was used to create projections of the refined map. Template-based automated particle picking in Gautomatch was then applied using 20 Å low pass filtered projections as templates. This picking was then applied on 7,266 movies with an estimated CTF resolution better than 6 Å. The new set of particles was pruned by 2D and 3D classification resulting in 37,619 particles being allocated to one class with well-defined features and apparent C2 symmetry. This class was further refined imposing C2 symmetry, and particles were corrected for beam-induced motion and CTF refined in RELION-3 (**Fig. S1**). A consensus map with nominal resolution of 3.3 Å based on the FSC curve at 0.143 criterion^47,48^ was obtained after post processing using an automatically estimated B-factor of −45 Å^2^ (**Fig. S3, S4**).

Particles considered in the consensus map were imported into cryoSPARC v2^49^ for localized refinement with the aim of improving quality of the densities in the "wing" and "foot" regions. Masks around these regions were created in UCSF Chimera^50^ using a local resolution filtered version of the consensus map as template and the volume segmentation tool. Both regions benefited from the local refinement procedure as the resulting maps display better connectivity and side chain densities, compared to the consensus map in the considered regions (**Fig. S2, S5**). Interestingly, the best local refinement maps of the wing region were obtained without performing any prior signal subtraction, as judged by visually inspecting the densities.

An atomic model of hTg based on the consensus, “foot” and “wing” maps was built in Coot^51^. The model covers 90% of the amino acid sequence, lacking mainly loops and the N- and C-terminus extensions. Maps were merged using the program phenix.combine_focused_maps and the atomic model was real space refined in PHENIX^52^ and validated using MolProbity^53^.

## Abbreviations

ChEL: Cholinesterase-like domain
Cryo-EM: Cryogenic transmission electron microscopy
CTF: Contrast transfer function
DIT: Diiodotyrosination
DSB: Disulfide bond
GlcNAc: N-linked acetylglucosamine
hTg: Human thyroglobulin
MIT: Monoiodotyrosination
MS: Mass spectrometry
NIH: Non-homologous insertions
PTM: Post translational modification
Tg: Thyroglobulin
TH: Thyroid hormone
T3: Triiodothyronine
T4: Thyroxine

## Accession codes

The EM map for the complete hTG molecule has been deposited in the EMDB under accession code EMD-12073. Atomic coordinates for hTG have been deposited in the Protein Data Bank under the accession code PDB 7B75.

## Acknowledgments

This work was funded by the Swiss National Science Foundation (NCCR TransCure). We thank K. Goldie and L. Kovaczik for support in electron microscopy. Calculations were performed at sciCORE (http://scicore.unibas.ch) scientific computing center at the University of Basel. The Novo Nordisk Foundation Center for Protein Research is supported financially by the Novo Nordisk Foundation (grant NNF14CC0001). N.M.I.T. is a member of the Integrative Structural Biology Cluster (ISBUC) at the University of Copenhagen.

## Notes

### Competing Interest Statement

The authors have declared no competing interest.

