## Supplementary Information for "Cryo-EM structure of native human thyroglobulin"

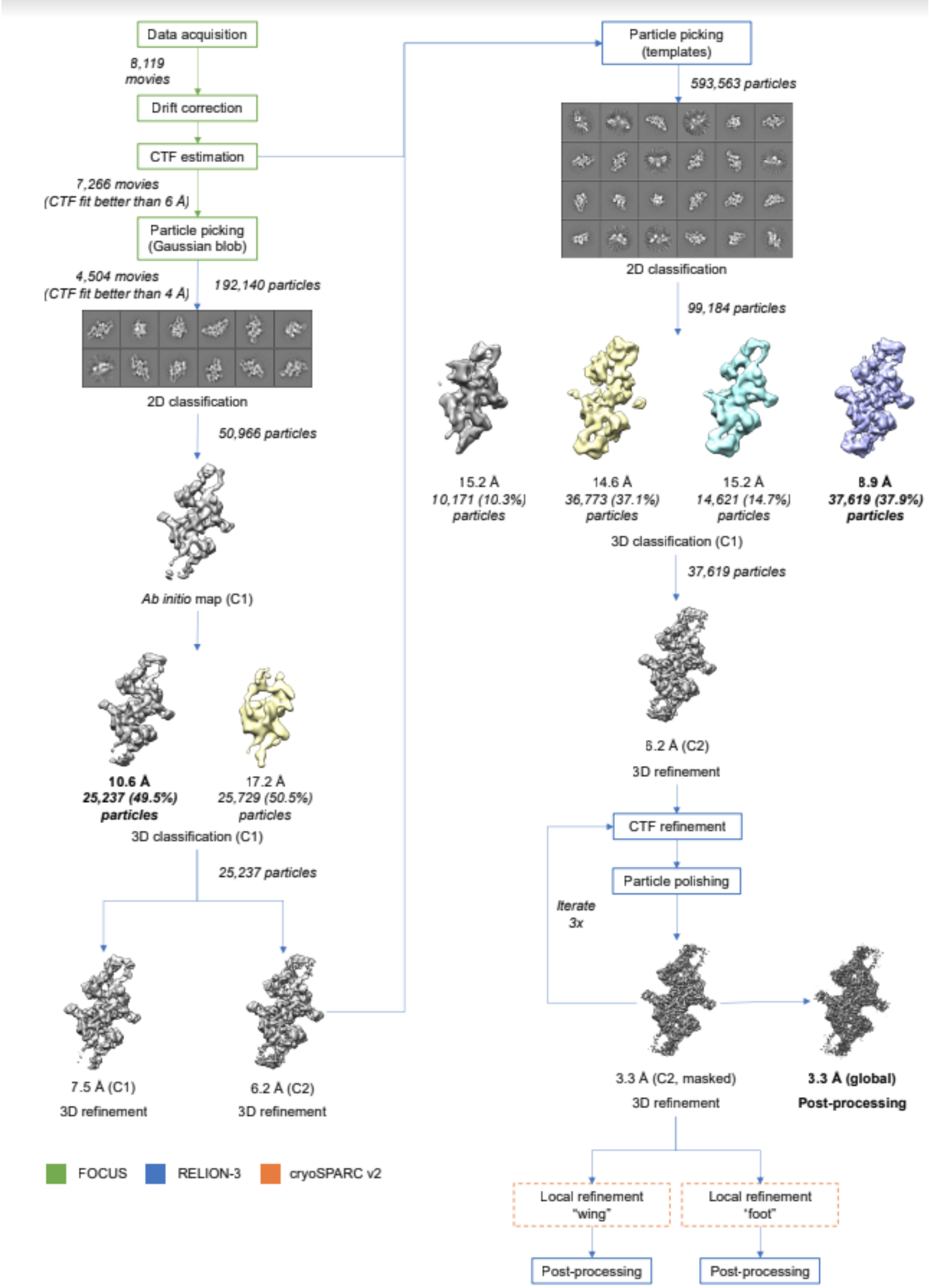

**Figure S1.** Processing workflow of hTg cryo-EM dataset.

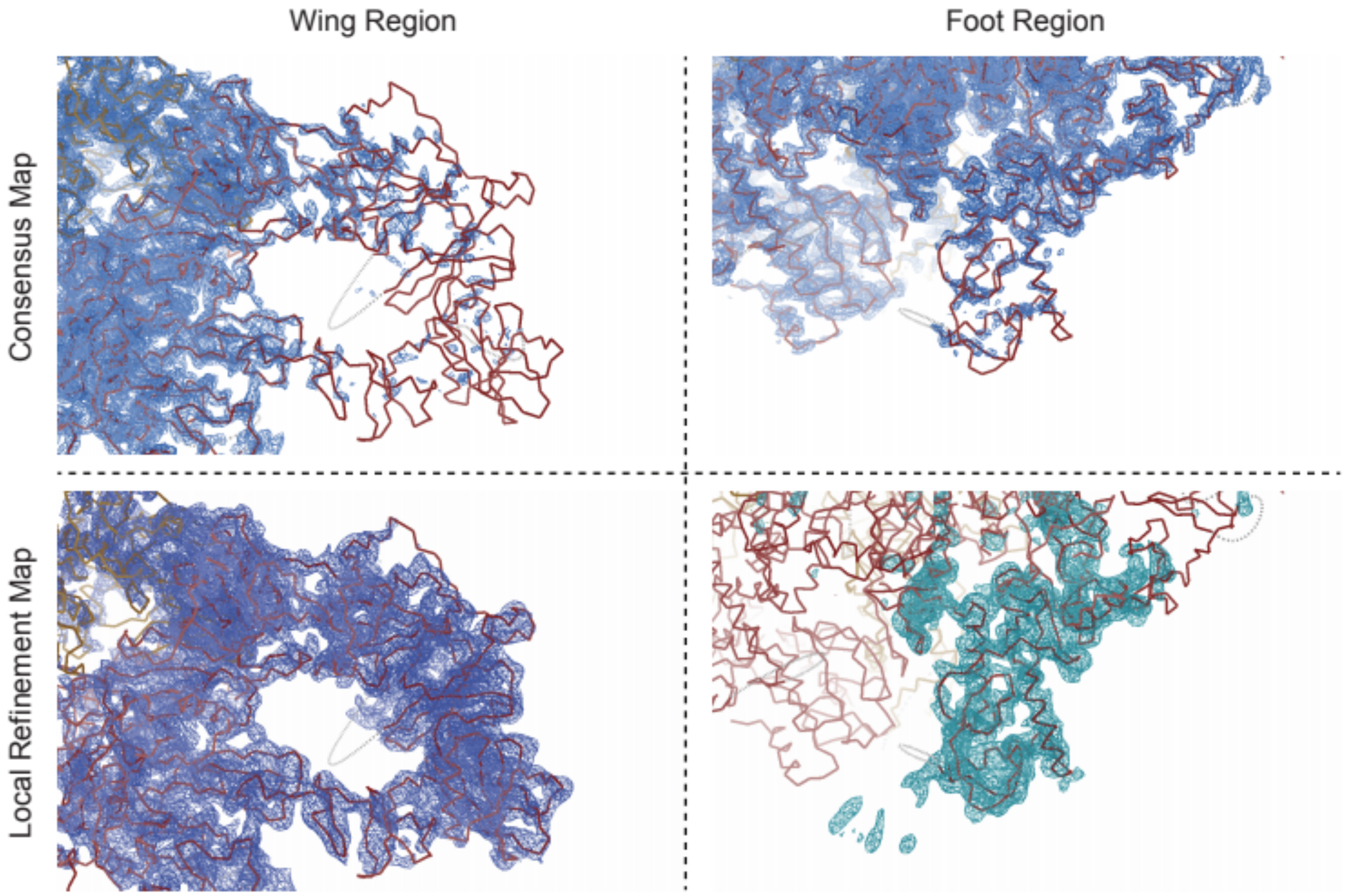


**Figure S2**. Comparison of the “foot” and “wing” regions in the consensus map and in the local refinement maps at equivalent threshold levels. Note the foot local refinement was performed with signal-subtracted particles while the wing was not.

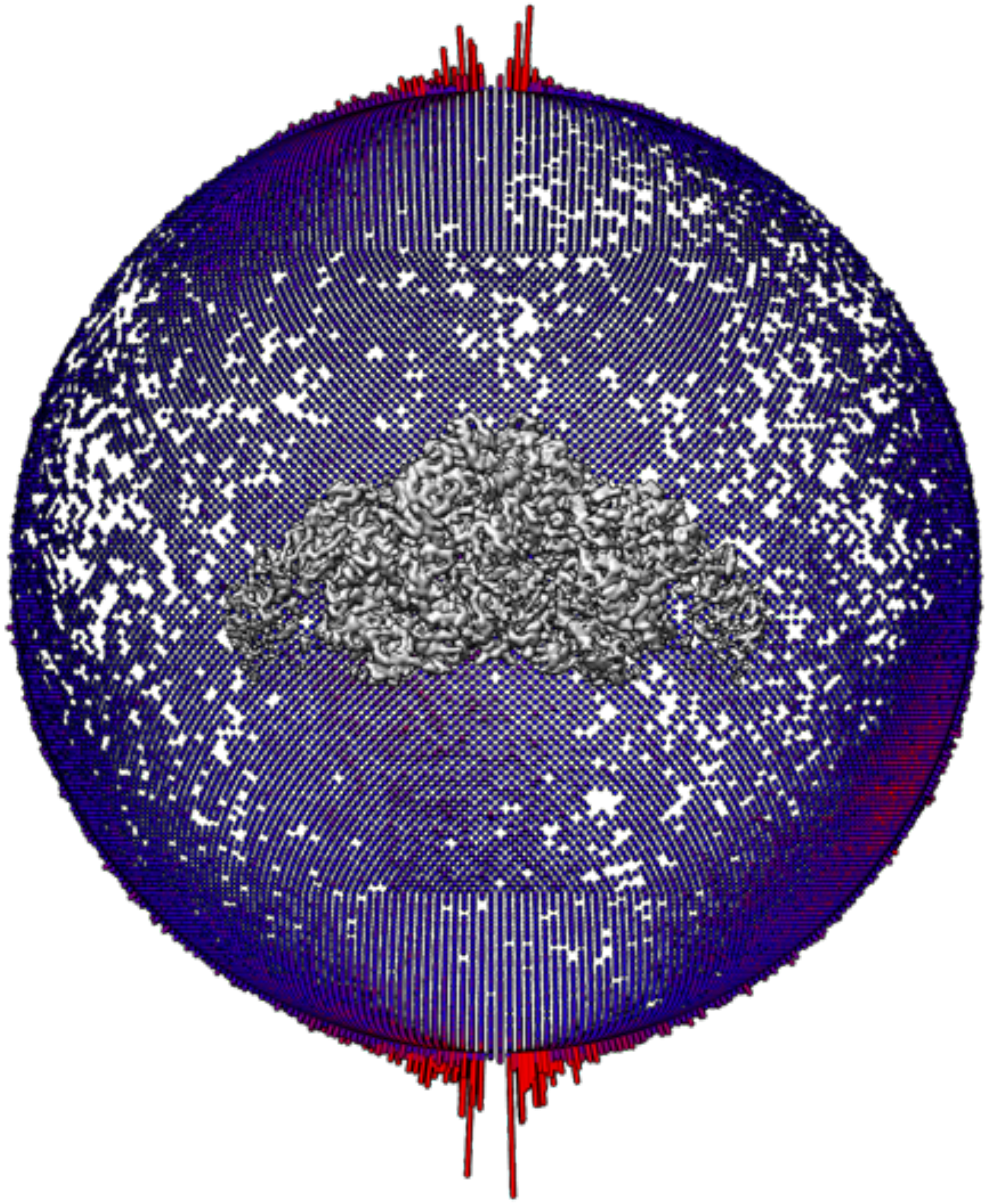

**Figure S3**. Angular plot distribution of the hTg consensus map.

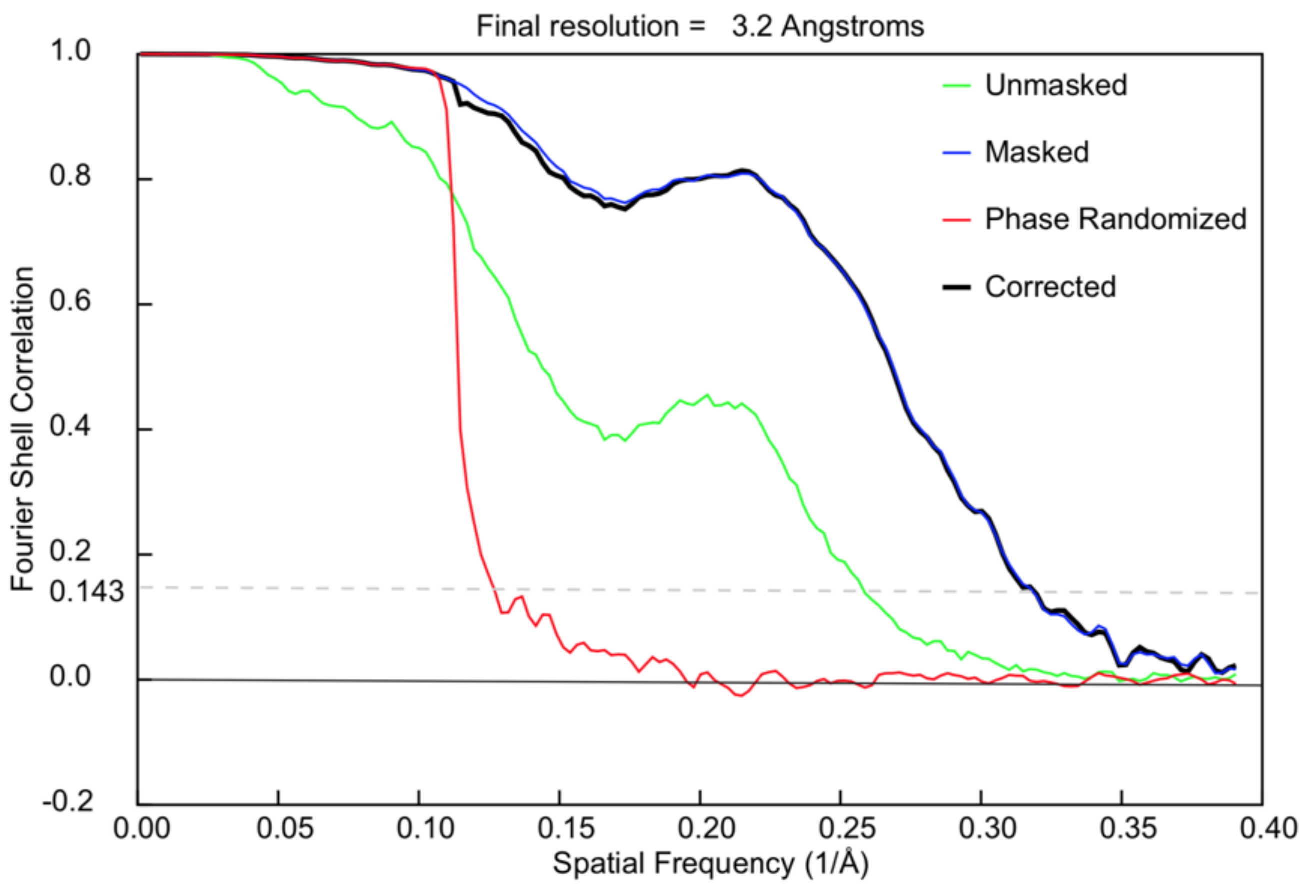


**Figure S4**. FSC plot of the hTg composite map.

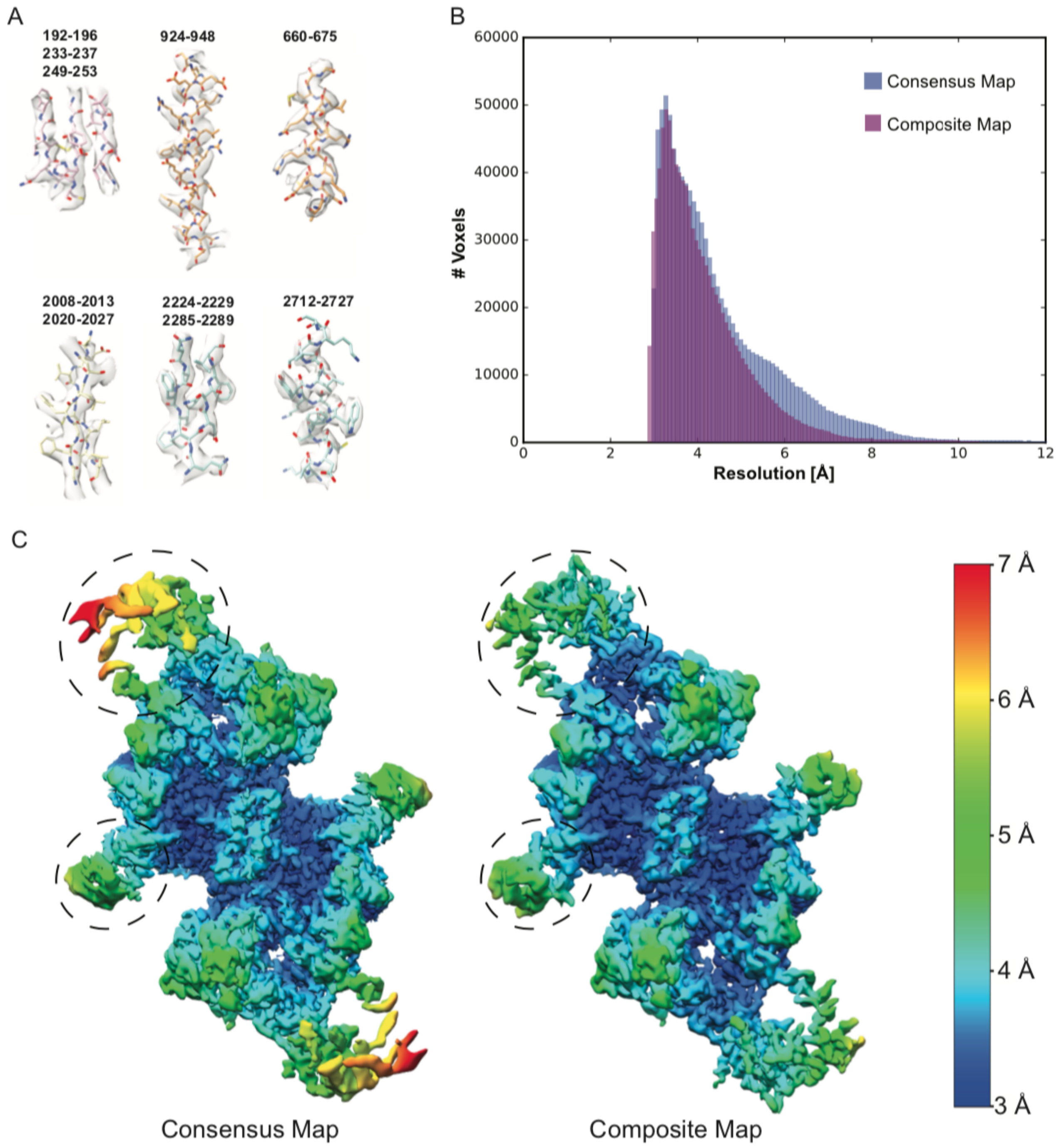


**Figure S5**. (A) Selected fragments of the refined hTg atomic model shown inside the composite electron density map. (B) Local resolution histograms of the consensus and composite hTg maps. (C) Consensus and composite electron density maps colored by local resolution with flexible “wing” and “foot” regions in dashed ovals.

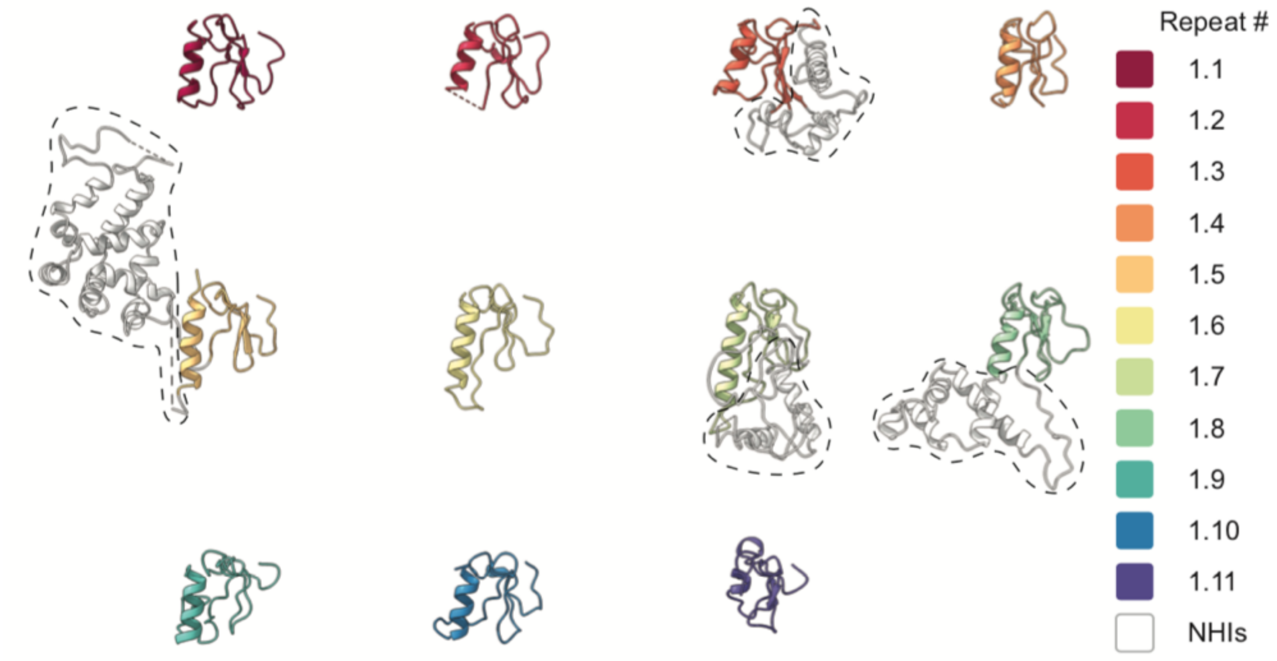


**Figure S6**. Type 1 repeats aligned. Color code for each repeat on the right. NHIs are highlighted with a dashed line.
